# Myth-busting the provider-user relationship for digital sequence information

**DOI:** 10.1101/2021.08.02.454535

**Authors:** Amber Hartman Scholz, Matthias Lange, Pia Habekost, Paul Oldham, Ibon Cancio, Guy Cochrane, Jens Freitag

## Abstract

The United Nations Convention on Biological Diversity (CBD) formally recognized the sovereign rights of nations over their biological diversity. Implicit within the treaty is the idea that mega-biodiverse countries will provide genetic resources and grant access to them and scientists in high-income countries will use these resources and share back benefits. However, little research has been conducted on how this framework is reflected in real-life scientific practice. Currently, parties to the CBD) are debating whether or not digital sequence information (DSI) should be regulated under a new benefit-sharing framework. At this critical time point in the upcoming international negotiations, we test the fundamental hypothesis of provision and use by looking at the global patterns of access and use in scientific publications. Our data reject the provider-user relationship and suggest far more complex information flow for digital sequence information. Therefore, any new policy decisions on digital sequence information should be aware of the high level of use of DSI across low- and middle-income countries and seek to preserve open access to this crucial common good.

## Background

The Convention on Biological Diversity is the international policy mechanism to reduce species, habitat, and ecosystem loss on this planet. The three overarching goals of the CBD, agreed upon in 1992, are conservation of biodiversity, sustainable use of this biodiversity, and fair and equitable benefit sharing from genetic resources. The third goal represents a political “balancing act” with the first two goals because it is intended to incentivize access and use of genetic resources (GR) so that benefits from use of biodiversity will flow back to the providing country and thus encourage conservation and support the first two goals.

Although it is officially recognized by parties to the CBD that all countries are both users and providers of GRs, in practice, most low- and middle-income countries (LMICs) see themselves predominantly as providers and, conversely, many high-income countries (HICs) view themselves as users [1]. While the CBD originally envisioned a facilitation mechanism for access to GR, the Nagoya Protocol (negotiated in 2010) codified a bilateral system in which a single country gives permission to a single user which has perpetuated the provider-user paradigm [2]. In fact, the complex legal landscape that has resulted from the post-2010 implementation of the Nagoya Protocol reflects this. HICs often focus on user compliance [3] and LMICs focus on access laws even though every country should theoretically be responsible for user checks [4]. (Countries are not bound by the Nagoya Protocol to regulate access.) For example, to our knowledge, to date only developed countries have implemented user compliance mechanisms (i.e. laws that check if users have complied) with provider country laws, most notably the European Union [5] and Japan [6].

However, whether patterns of scientific use of GR actually follow these user-provider assumptions is not a question that has received much attention [7]. GR provision and use is difficult to follow since GR sampling and exchanges are not centrally administered or recorded. However, the use and citation of sequence data from GR in scientific publications enables a “proxy” view on provider-user relationships and happens to be itself highly relevant to the current CBD discussions.

In CBD policy circles, nucleotide sequence data (as well as potentially other data types) are known as “digital sequence information” (DSI) [8]. Because of the exponential growth and widespread use and reliance on DSI in the biological sciences, the political question of the hour is whether and how benefit-sharing from DSI should be required. Parties will decide at the 15th Conference of the Parties, which has been delayed by the pandemic but tentatively scheduled for October 2021, whether DSI should be treated like GR, whether monetary and/or non-monetary access and benefit-sharing will be required and documented, and, if so, whether the policy framework for benefit-sharing will be bilateral or multilateral [9]. Thus, COP15 will be an important milestone for policymakers and scientists alike, making the question of patterns of use of DSI, presented here, quite timely.

Because many negotiators at the COP15 will be familiar with the bilateral mechanisms of the CBD and its Nagoya Protocol, it is likely that the default pre-conception around DSI for most negotiators will be the “provider-user dichotomy” assuming a primarily uni-directional (roughly global south to north) provision and use relationship. This is actually a hypothesis that can be tested with data from open access public DSI databases, in which the country of origin for the DSI can be found, and via publication databases, where use of DSI can be assessed by proxy through the affiliations of the authors which can be parsed into geographical locations. While keeping in mind the potential shortcomings and accuracy issues [10], here we test this hypothesis and display the results in a free and open data analysis platform with the aim of analyzing whether a real directionality exists from provider country to DSI user country, with LMICs in one side and HICs on the other. The data and their implications will hopefully lead towards informed decisions and evidence-based policymaking.

## Data Description

This article is a companion paper to a data note submitted in the same issue of this journal which was made available as a pre-print in BioRxiv [11]. A web application is provided to explore and visualise the dataset at http://wildsi.ipk-gatersleben.de and, for clarity, we have compiled here a brief excerpt of the dataset described in the data note.

The vast majority of scientific journals and most funding agencies require that DSI be made freely available, at the latest, by the time of publication. Submissions of sequence data are required by journals as a condition of publication and rely upon the use of unique identifier(s) (called accession numbers, ANs) generated by a member of the International Nucleotide Sequence Database Collaboration (INSDC). During the sequence submission process, metadata associated with the DSI is also submitted including, where appropriate, the country field (data field “/country) which is defined as “locality of isolation of the sequenced sample indicated in terms of political names for nations, oceans or seas, followed by regions and localities.”

At the time of these analyses, there were 17,816,729 sequences in the INSDC with a country tag. Generally, sequences with country information come from a natural environment. For this study, we did not perform subsequent analyses on the taxonomic distribution of these sequences, which was previously assessed in [12]. Access to GR is needed to produce DSI. In this paper we use the term “provider” to designate the /country information found in the INSDC and indicate the geographical location from where the GR, and thus indirectly the DSI, originated. (Note that “provider” does not reflect where the sequencing was done or the entity that made the research/funding investment.)

For each of the >17.8 million ANs, if a publication was listed in the sequence entry page (within the INSDC database), this was added to the dataset as a “primary” publication. In a parallel step, the European PubMedCentral (ePMC) open-access publication database was text-mined for all >17.8 million sequence ANs. If a publication listed any of these sequences, it was added to the dataset as a “secondary publication”. A total of 117,483 primary and/or secondary publications were included in this analysis. Publications citing the use of DSI are representative of DSI scientific “use”. The associated author metadata from the primary and secondary publications was machine read and parsed. The geographical location of the first author was identified where data quality was sufficient. We note that first author information presents a restricted view of author networks that reflects limitations in the availability of full author information. As more author data becomes available, we anticipate that it will be possible to engage in more detailed analysis of author networks. This dataset forms the basis of the DSI “user” geographical locations. Additional quality control, data parsing, table merging, and data visualization steps were required that are further explained in the corresponding data note [11]. We make no further classification under the term “use”, as our methods at this stage cannot distinguish among the different types of use of DSI, for example, commercial versus non-commercial. On average, we expect that many peer-reviewed publications are more likely to derive from non-commercial research.

## Analyses

The first question addressed is which countries are currently providing DSI to the global dataset available through the INSDC (Figure 1). The largest providers of DSI are currently not LMICs but are the United States, China, Canada, and Japan, providing roughly half of the global dataset, although large middle-income countries such as India and Brazil are in the next wave of providers (Fig. 1a). In the two years since Rohden et al. [12] described this trend for a CBD-commissioned study the pattern has not significantly changed. Although, it is important to note that only 14.6% of all sequences in INSDC had country information available. This is down slightly from the 16% observed in the April 2019 global sequence dataset analyzed by Rohden et al. However, this does not necessarily represent a statistical trend. There are multiple factors that could cause this seeming decrease. For example, large deposits of sequences that are not appropriate for country labelling, e.g., human data could grow the dataset but decrease the amount of country-labelled DSI. This was not further investigated.

**Figure 1.**
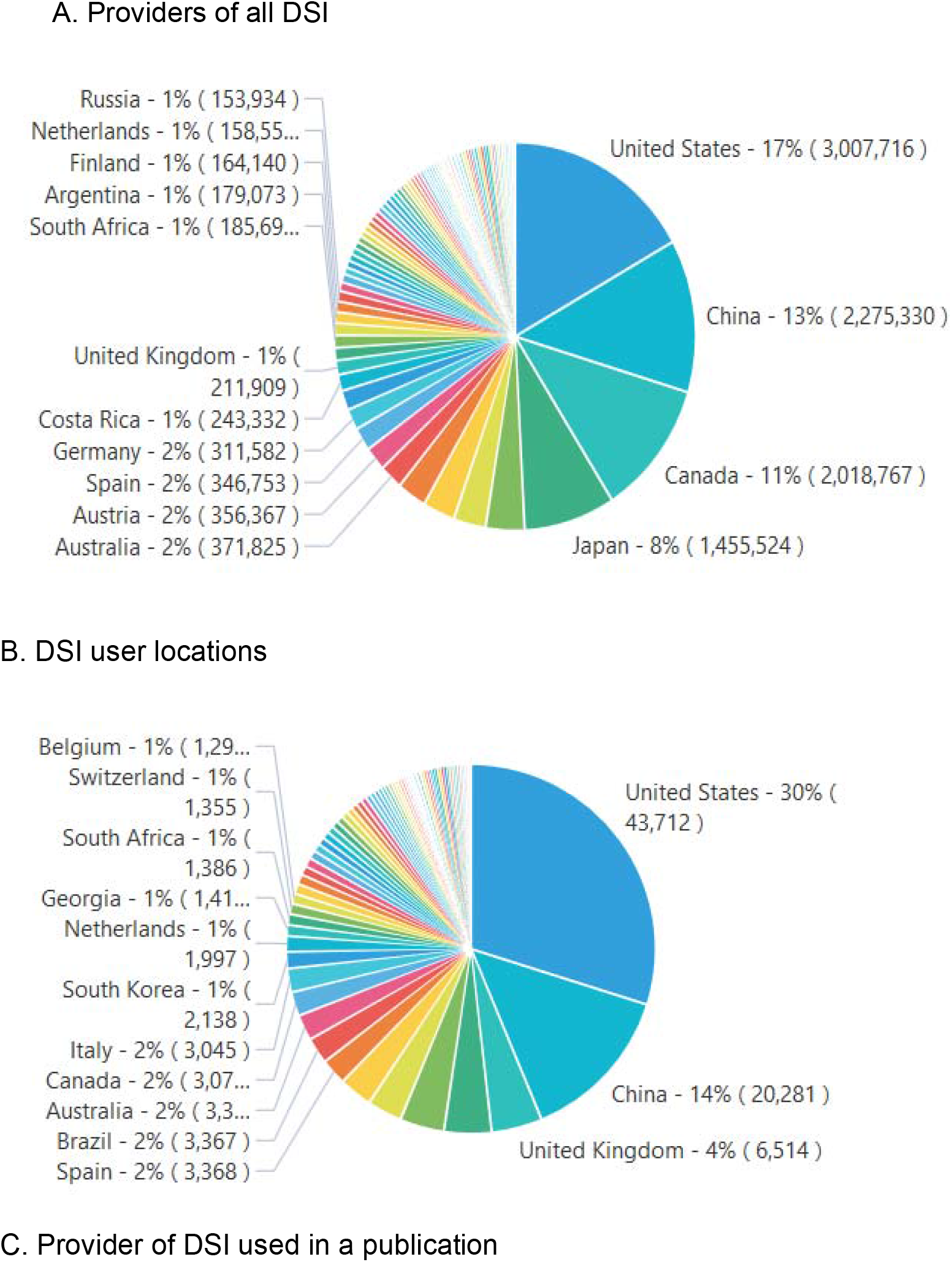
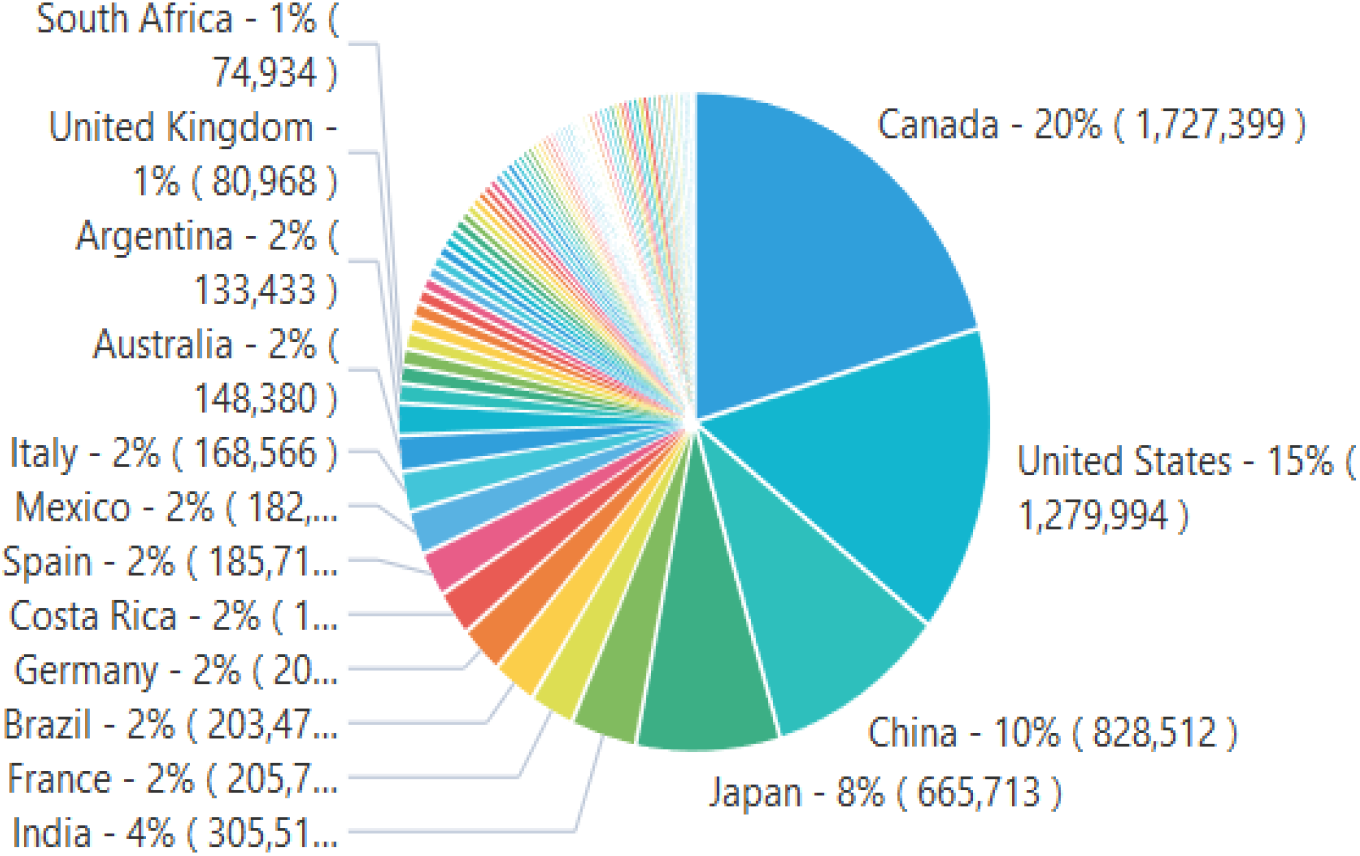
a) Countries that “provide” (that is, are the countries of origin) for DSI. b) the geographical location of users (authors in scientific publications that reference DSI), c) the geographical distribution of the provider countries of the DSI cited in 1b. Interactive charts are available on the web platform under “1. Overview of DSI use”.

Once these DSI are made available by provider countries, the natural question is to ask “who” is using the DSI, i.e., scientists sitting in which countries are publishing (which we call “using”) and citing DSI in a publication (Fig. 1b). In order to begin to understand the provider-user relationship, it is also important to understand where the DSI they are citing comes from, i.e., which countries provided acces to the GR (Fig.1c). To summarize, most DSI is being provided and used by HICs. Although there is DSI use and provision in many different directions. These data do not support the idea of a uni-directional provider-user dichotomy.

To further understand the real-world provider-user relationship, another angle to examine is the relationship between a country’s use of its own “national DSI” (where it was the provider country of the GR) versus its use of DSI from other countries: We call this the “self:world” ratio where self-use of DSI is divided by use of DSI from all other countries (Figure 2).

**Figure 2:**
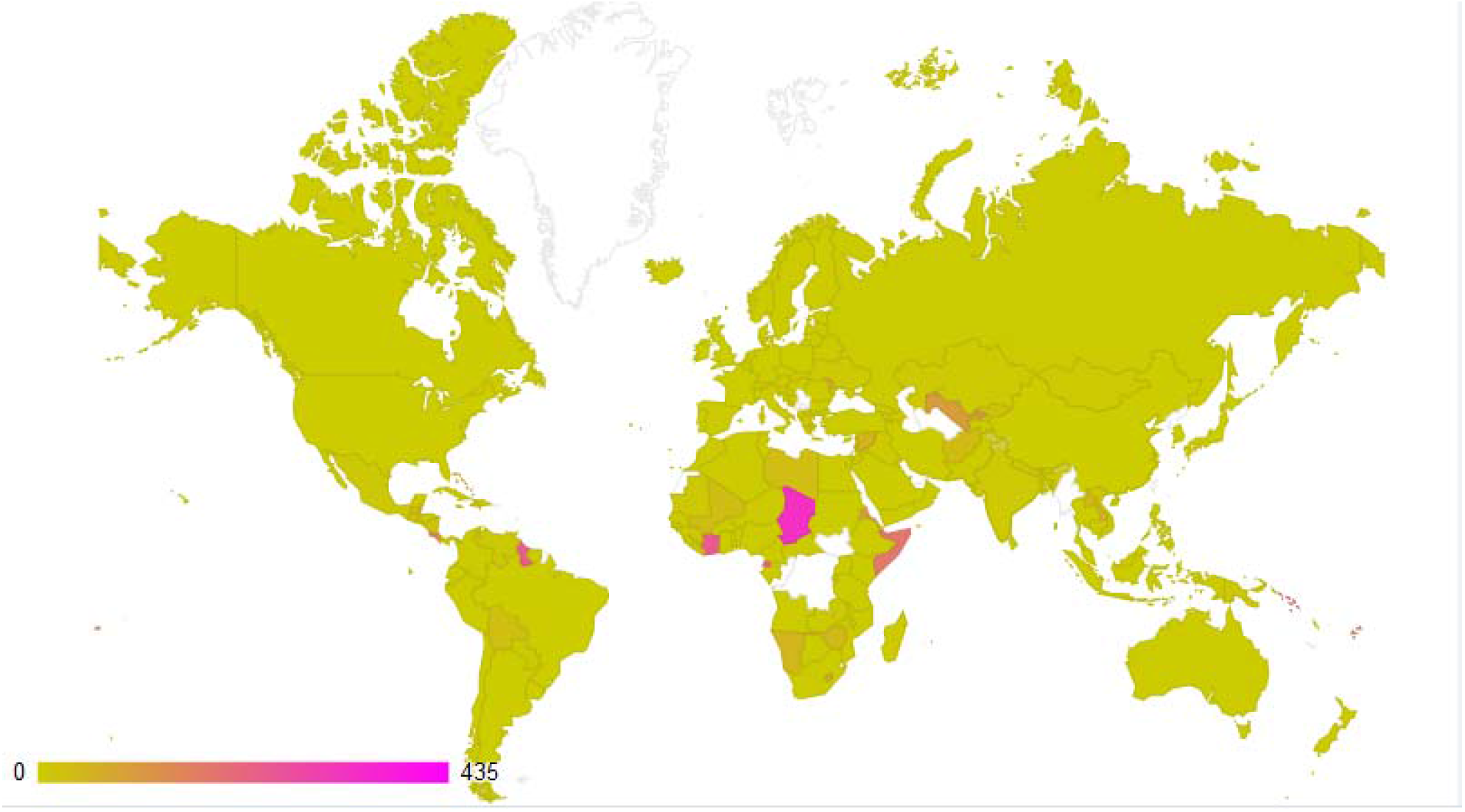
This is the ratio between the foreign use of a given country’s DSI (nominator) and the domestic use of a country’s DSI. Green is a balanced use of DSI between foreign and domestic DSI. Pink is an indication of strong foreign use of DSI with less domestic use. An interactive chart is available on the web platform under graph 3.4.

Although there are significant differences in the number of scientists and the volume of use and re-use between LMICs and HICs, the relationship between a country’s use of its own DSI and use of the global DSI dataset (all non-national DSI) is relatively homogenous. A few countries have more “foreign use”, i.e. their national scientists use less relative to non-national scientists, such as Chad and Guyana. However, the vast swaths of green suggest that the ratio between national and foreign use of DSI is relatively even.

At international meetings, including the CBD Conference of the Parties, countries can form negotiating blocs that enable coordination between similar perspectives and sharing of preparation work when developing negotiating positions. These blocs often represent underlying economic similarities between countries. To understand broad trends through a similar lens used in these political discussions, we grouped countries into three overarching groups: low-income countries in a group called G-77, middle-income countries known as BRICS (Brazil, Russia, India, China, South Africa), and high-income countries under OECD (Organization of Economic Cooperation and Development). These broad groupings, although imperfect, allow for visual representations that could proxy common trends within UN political discussions.

Three significant trends can be observed in figure 3. First, the largest color block of users (counted by a publication not an individual) in each bar corresponds with the economic bloc of the country of origin of the DSI. For example, the biggest group of users using G77 DSI is G77 users. This suggests that users tend to use DSI from their own country/bloc rather than from outside. They research and publish more with “locally-sourced” DSI. This data again rejects the unidirectional provider-use hypothesis. If that hypothesis were true, then OECD users (from HICs) should be the biggest bloc in all three bars.

**Figure 3:**
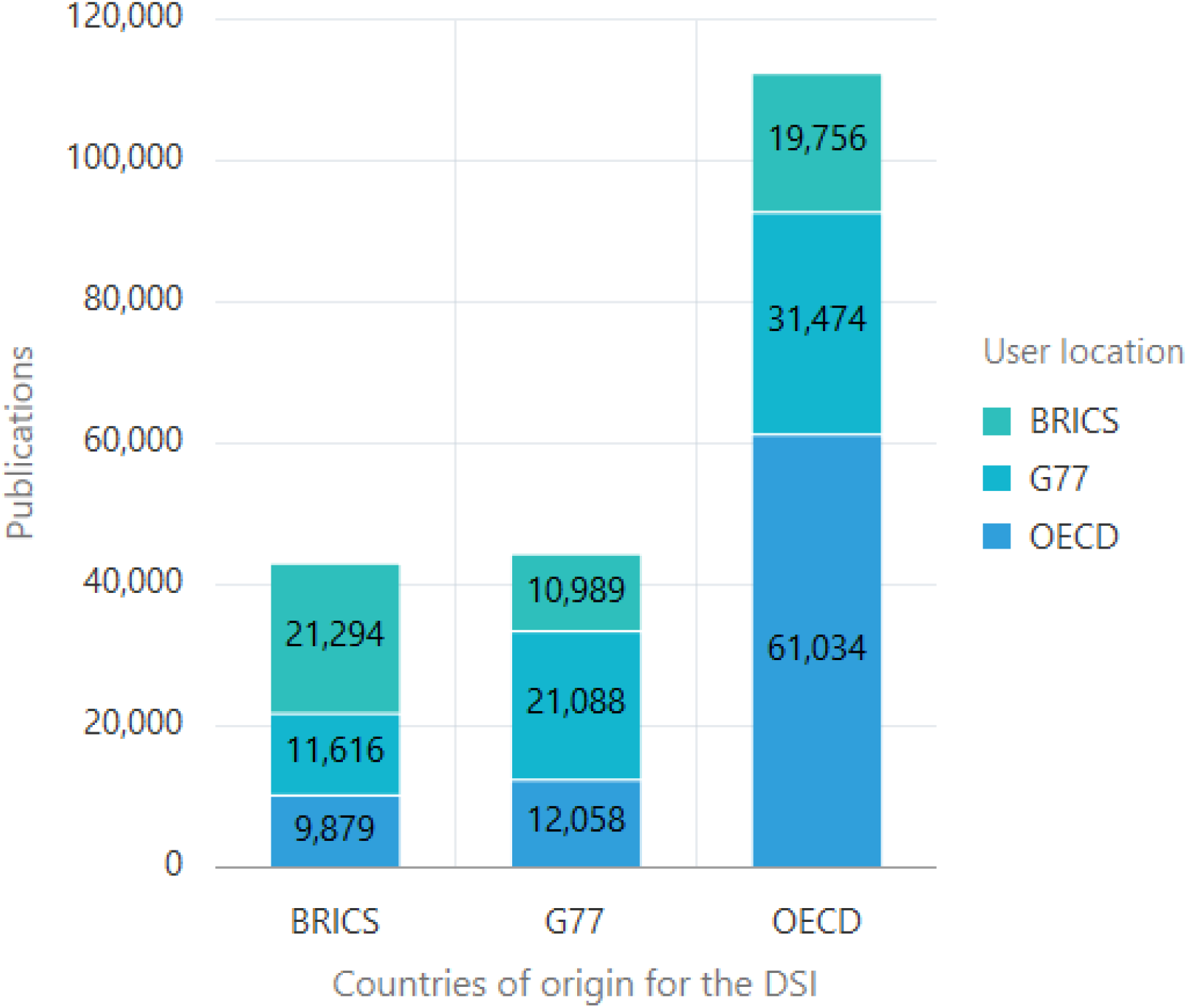
DSI use by economic blocs. On the x-axis is the country of origin of the DSI (/country field in the sequence database). On the y-axis are the total number of publications. The color blocks represent the geographical location of the users (authors). An interactive chart is available on the web platform under graph 5.1.

Second, OECD DSI is used nearly 3 times more in comparison to both BRICS-sourced DSI and G77-sourced DSI. This is shown by the difference in the height of the bars. This is likely due mostly to the fact that there is simply a lot more OECD-sourced DSI in the databases than DSI from the other blocs. Many negotiators feel strongly that DSI sourced from mega-biodiverse countries is inherently more usable and valuable than DSI from other sources. The data shown here do not support this, otherwise G77 or BRICS DSI would be used more than OECD DSI. Finally, the graph also shows that there are important gaps between the different blocs. There are fewer total DSI-related publications coming from authors in G77 and BRICS than from OECD-based authors (roughly 30-40% fewer). The use of DSI and, likely biological research in general, has lower total output as compared to OECD countries. However, the scale of the data also shows that G77 and BRICS-based authors are still quite scientifically productive. We suspect, but cannot presently confirm, that a fuller view of the author landscape would further emphasize the contribution of G77 and BRICS-based authors.

The fourth question we were able to investigate with this dataset is the geographical interconnectedness between DSI researchers. To this end, two network diagrams were built. The “providing” network displays which countries are using a given country’s DSI (i.e., the countries to which country X is providing”). And the “using” network displays the countries whose DSI are being used by country X’s scientists. These data are also helpful to show that both neighboring and distant countries use DSI from many countries.

In Figure 4, DSI provisioning and use for Malaysia, which is both a G77 member and a mega-biodiverse country, is shown as an example. In Figure 4a, many LMICs (and not just HICs) use DSI from Malaysia, for example, Zambia, India, Peru, and Mexico to name a few. Conversely, in Figure 4b, scientists in Malaysia use DSI from 68 countries. Again, here there is no evidence of a provider-use relationship in DSI usage. Rather, Malaysian scientists use (cite in publications) DSI from a wide variety of countries and economic settins including Germany, Norway, Costa Rica, and Ghana to name a few. This data complements the data presented in Figure 3a which suggests that, although scientists use their own national DSI more frequently, when they use foreign DSI they do not appear to be primarily using DSI from biodiversity-rich countries but rather DSI from across the world without any clear geographical or economic clustering patterns.

**Figure 4.**
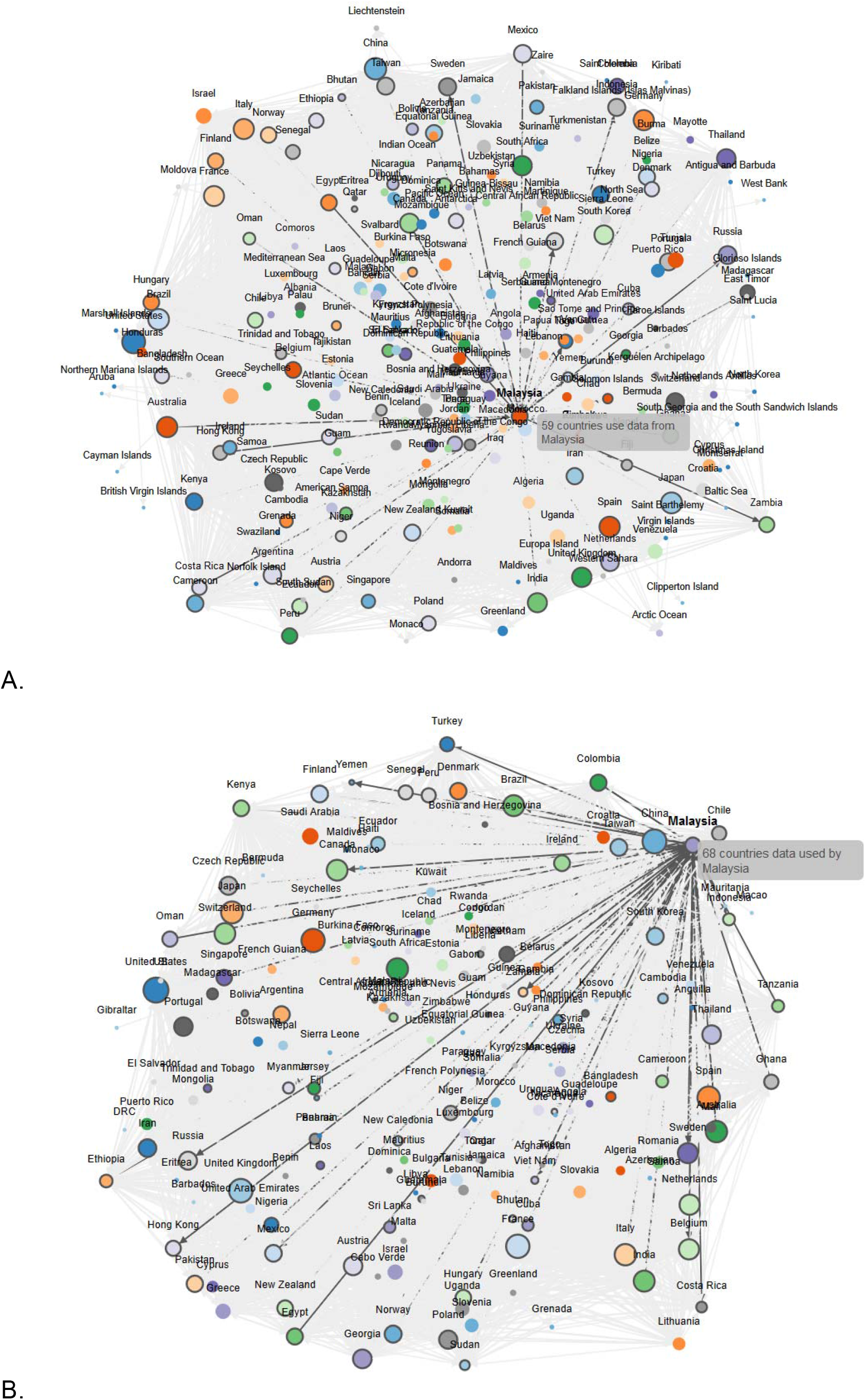
Networks diagrams displaying country-based DSI provision and use patterns. In both graphs, data from Malaysia was selected as an example. a) 59 countries are using data from Malaysia. b) Malaysian scientists are using DSI from 68 countries. An interactive chart is available on the web platform under graph 6.1 and 6.3. Note: Neither the length of the connecting arrows nor the clustering reflects a statistical relationship or indicates stronger linkage between connected countries because the clustering algorithm is random distribution.

A final question that can be addressed is what the overall providing-use relationship is for every country and whether there is a global trend to this relationship. Indeed, given the linear trend displayed in figure 5, it seems that many countries provide and use DSI from a roughly equal number of countries. In other words, if scientists from a given country are providing DSI, they are often using DSI at a similar level. However, for small countries, especially LMICS, with accordingly smaller datasets and scientific communities, tend to cluster in the bottom left of the graph meaning they have little provisioning of DSI and even less scientific use of DSI.

**Figure 5.**
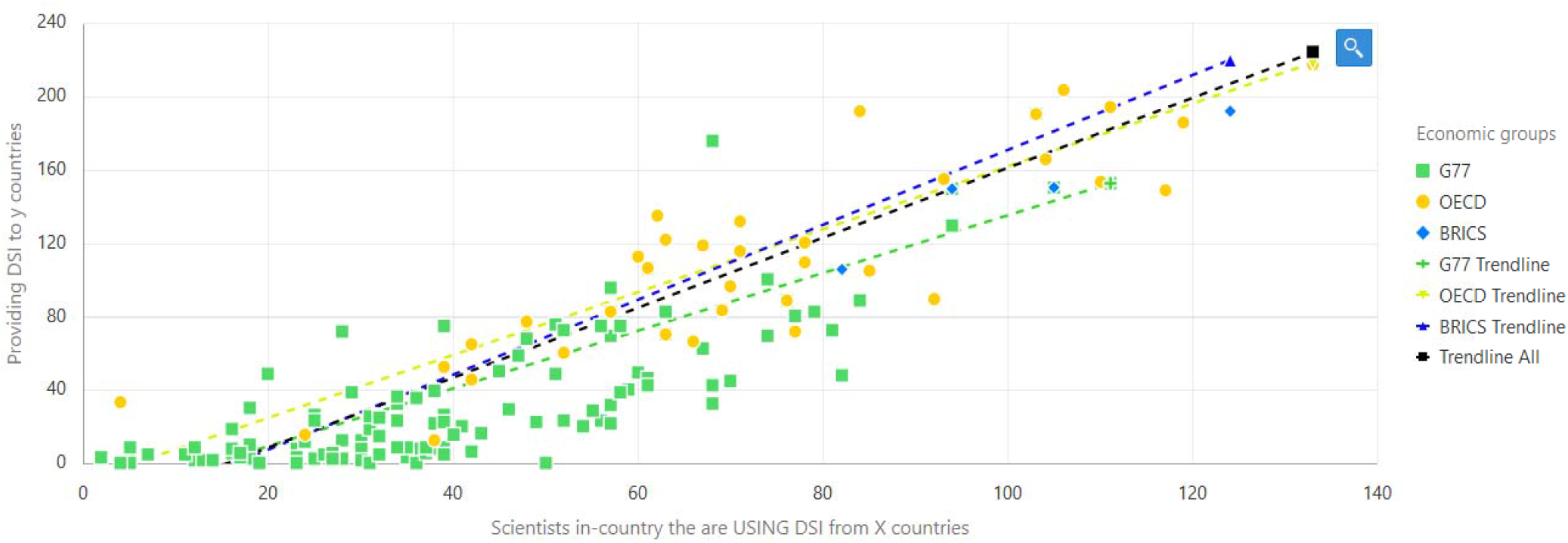
Relationship between use and provision of DSI. The x-axis displays the use of DSI by a country’s scientists and the y-axis displays the provisioning of DSI by a given country. An interactive chart is available on the web platform under graph 6.3.

## Discussion

Preconceptions in policymaking can be tested by looking at empirical data. Because there is a central repository for DSI, data and global analyses can be conducted to inform the debate around benefit-sharing and test hypotheses. The data presented here show that the concept of a user-provider dichotomy bias for DSI is rejected.

This suggests that if an ABS policy mechanism for DSI incorrectly assumes a uni-directional provider-user relationship, this could negatively impact scientists in LMICs the hardest. Scientists in LMICs are often resource-limited and have more personnel and infrastructure constraints than scientists working in HICs. Given current political discussions, LMIC-sourced DSI would be the most likely to fall under an ABS regime, but it is LMIC researchers that are the predominant users of this DSI (Figure 3). If DSI policymakers do not recognize the “self-use” of DSI, that is the use of that country’s own DSI by in-country scientists, then they could potentially do great harm to scientists in LMICs which could have long-term implications for their domestic bioeconomy strategies and broader research and innovation goals. There are indeed important inequalities across the globe, but a DSI ABS system should try to reduce these inequalities rather than exacerbate them.

The global goal, in our view, ultimately should be to increase the scientific output and generation of DSI from G77 and BRICS countries to similar levels as those observed within the OECD and shown in Figure 3. Increased research capacity in LMICs would benefit everybody globally and global biodiversity knowledge gaps, including those identified by the Global Biodiversity Framework, could be better covered. In order to do this, any DSI policy mechanism should recognize the existing divide and encourage DSI use, publication, and collaboration, perhaps explicitly dedicating significant capacity building to scientifically leveling the DSI playing field. This would be a much different approach than a “lock-it-up and control it” approach to DSI, which would negatively affect researchers everywhere on the globe and especially those in LMICs.

Furthermore, provisioning, use, and re-use of DSI as interpreted through DSI citation in publications is not a one-way street but rather a multi-directional traffic circle of data flowing in many different directions amongst all countries of the world. DSI is used by neighboring countries and distant countries, by LMICs and HICs alike without any clear patterns, regional, or economic trends (Figure 4). Additionally, the network diagrams remind us that scientists working in developing countries as well as the biological diversity of developed countries are often overlooked in political discussions that oversimplify provision and use of biological data.

The one clear trend observed in this analysis is that providing and using tend to go hand in hand (Figure 2 and 5). Large countries tend to use and provide a relatively large amount of DSI and smaller countries use and provide less but trends based on development status (or, indirectly, the presence mega-biodiversity) were not detected. In general, the relationship between use and provision seems to be relatively linear and not biased towards HICs but, instead, slightly biased towards LMICs (see LMIC trend line, Fig.5).

The dataset presented here is not a comprehensive dataset of all publications citing DSI but is, in fact, limited to the open-access publications available for text-mining in ePMC as well as other dataset limitations explained in [11]. Furthermore, we note that the dataset would be further improved if the country of origin information (14.6% of sequences in this dataset had such information) provided by scientists submitting sequence data to INSDC were more consistent and compliance with the requirement to submit this information for relevant DSI were bolstered. With this information, clearer references to regional conditions would be possible and thus more valid scientific statements and analyses would be possible. For example, gene-function relationships could be mapped more precisely with climatic, geological or atmospheric features.

However, acknowledging the above limitations, this analysis is the largest and only comparison of this size and perspective to-date and represents a novel attempt to bring data and a new perspective on DSI to the policymaking process. We encourage other groups to expand and build upon this dataset for other policy environments and to re-use and complement these data with additional perspectives. Future studies are planned that will expand this dataset to include closed-access publications and the patent system [13], where greater insights on potential commercial use of DSI can be assessed. Furthermore, assessments are planned that will provide first analyses on the field of study and taxonomic patterns.

## Potential Implications

These data also raise a fundamental question about current ABS frameworks already in place for GR, especially the Nagoya Protocol. For GR there is no central repository for movement across national borders, but these data suggest that the provider-user relationship for GR could follow similar patterns to those observed for DSI. If so, this could suggest the existing bilateral system and the predominance of user checks in HICs (rather than globally) is perhaps not the most appropriate way to ensure benefit-sharing. While provocative, this could suggest that policymakers, in the future, ought to revisit ABS frameworks from the bottom up.

Future political decisions around how to handle DSI should account for the complexity of geographical provision and use trends presented here. Policymakers need to appreciate the tremendous contribution towards non-monetary benefit sharing that these global biological and publication databases make towards broader CBD goals and towards the SDGs. As policymakers open COP15 in October and meet in person in May 2022 to finalize a decision on DSI and the Global Biodiversity Framework, we hope these data will make a constructive contribution to an evidence based DSI policymaking process.

## Methods

The methods used in this article are described in the companion paper which is a data note in the same issue of this journal also available as a pre-print in BioRxiv [11].

## Data availability

The dataset, figures, supplemental figures, and web application are available at http://wildsi.ipk-gatersleben.de.

### Abbreviations

ABS: access and benefit sharing
AN: accession number
BRICS: Brazil, Russia, India, China, South Africa
CBD: Convention on Biological Diversity
COP15: 15^th^ Conference of the Parties to the Convention on Biological Diversity
DSI: digital sequence information
ePMC: European PubMed Central
G77: the Group of 77 representing mostly low-income countries
GR: genetic resources
HICs: high-income countries
INSDC: International Nucleotide Database Collaboration
LMICs: low- and middle-income countries
OECD: Organization of Economic Cooperation and Development
SDGs: Sustainable Development Goals
UN: United Nations

## Competing Interests

The authors declare that they have no competing interests.

## Funding

This publication was made possible by the research project WiLDSI (Wissenschaftliche <u>L</u>ösungsansätze für <u>D</u>igitale <u>S</u>equenzinformation) funded by the German Federal Ministry of Education and Research (BMBF) under funding code 031B0862.

## Author’s contributions

AHS wrote the manuscript. AHS, GC, JF, and ML designed the research with critical input from IC and PO. ML and PH designed and programmed the data platform. All authors reviewed and edited the manuscript.

## Acknowledgements

We gratefully acknowledge the important technical input of our co-authors, Upneet Hillebrand, Mehmood Ghaffar, Blaise Alako, and Florian Zunder, in the related data note publication [11]. Their work enabled this focused policy analysis.

The authors

## Notes

http://wildsi.ipk-gatersleben.de

### Competing Interest Statement

The authors have declared no competing interest.

